# A genetically encoded marker for imaging membranes of *Plasmodium*

**DOI:** 10.1101/2025.10.24.684113

**Authors:** Leah Zink, Viola Reuschenbach, Samy Sid Ahmed, Muriel Lauer, Jana Niethammer, Aiste Kudulyte, Katharina Zorko, Oliver T. Fackler, Franziska Hentzschel, Markus Ganter

## Abstract

Fluorescence microscopy is a powerful tool to analyze subcellular architecture, and long-term live-cell imaging permits the analysis of the dynamics of intracellular structures. Applying this approach to the membranes of *Plasmodium* spp., the causative agent of malaria, yielded important insight. Fluorescent labelling of *Plasmodium*’s plasma membrane via membrane-resident proteins has been described, but proteins, which broadly mark internal membranes are currently elusive. Alternatively, general membrane dyes can be employed to study membrane dynamics, however, we find that the membrane dye BODIPY TR Ceramide has adverse effects on cell viability during live-cell imaging. To overcome this limitation, we present FLUMMI (Fixed-and-Live-cell Universal Membrane Marker for Imaging), a 32-amino acid long peptide derived from *P. falciparum* PCNA1, which targets (fluorescent) protein tags to internal membranes for detection in live or fixed samples. Importantly, FLUMMI enables non-invasive, long-term live-cell imaging of internal membranes without affecting *P. falciparum* viability and is functional in blood stages of both, the human malaria parasite *P. falciparum* and the rodent-infecting parasite *P. berghei*, as well as in *P. berghei* mosquito stages. FLUMMI is also compatible with advanced techniques such as expansion microscopy. Together, our results establish FLUMMI as a versatile and non-toxic tool for membrane imaging in *Plasmodium*.

## Introduction

Malaria remains a major public health burden with an estimated 263 million cases in 2023. Of the nearly 600,000 malaria-related deaths in the same year, most were children under the age of five in Sub-Saharan Africa (WHO, 2024). Malaria is caused by unicellular eukaryotes of the genus *Plasmodium*, with at least five species infecting humans. *P. falciparum* is responsible for the most virulent form of human malaria, which also accounts for the vast majority of cases and fatalities (WHO, 2024). To ultimately tackle malaria, a better understanding of the basic biology of *Plasmodium* spp. is needed.

A powerful tool to study *Plasmodium* spp. biology is fluorescence microscopy, which gives insight into the parasites’ subcellular architecture. Long-term live-cell imaging permits the analysis of the dynamics of internal structures such as organelles and membranes. For this, detectable marker proteins or specific dyes can be employed.

To visualize the parasite plasma membrane (PPM) in the rodent-infecting *P. berghei*, a reporter system based on the Plasma Membrane Protein 1 (PBANKA_0809400) was previously established (Burda *et al*., 2017). Similarly, the PPM of the human pathogen *P. falciparum* can be visualized, e.g., by endogenous tagging of the plasma membrane resident transporter 1 (PMRT1) (Wichers *et al*., 2022). However, similar genetically encoded fluorescent markers are currently lacking for the labeling of internal parasite membranes.

To mark all membranes independent of the expression dynamics and localization of a given reporter, commercially available BODIPY Ceramide dyes are commonly used. They combine the fluorescence properties of BODIPY with the sphingolipid ceramide, which supports membrane localization (Karolin *et al*., 1994; Marks *et al*., 2008). These dyes have been used to stain lipids in different *Plasmodium* species for live-cell imaging as well as fixed-cell imaging (Grüring *et al*., 2011; Liffner and Absalon, 2021; Liffner *et al*., 2023).

Despite the advantage of being a readily applicable and robust lipid dye, BODIPY Ceramide-associated phototoxicity during live-cell imaging was previously described (Kähärä *et al*., 2022), which is likely due to the formation of highly reactive singlet oxygen species close to the membrane (Liang *et al*., 2020). Thus, in particular for long-term live-cell imaging of *Plasmodium* spp., an alternative membrane marker may be useful.

To investigate the nucleocytoplasmic transport of *P. falciparum* PCNA1 (PF3D7_1361900), which transiently accumulates in a subset of nuclei of a *P. falciparum* schizont (Klaus *et al*., 2022), we predicted possible nuclear localization signals (NLS) and nuclear export signals (NES) in PCNA1 (Kosugi *et al*., 2009; Xu *et al*., 2015). To assess whether these predicted signals are sufficient for nuclear or cytoplasmic targeting, we fused them to GFP. We serendipitously found that a 32-amino acid predicted nuclear localization signal of *P. falciparum* PCNA1 targets protein tags to membranes, and here developed this peptide into a membrane marker for imaging.

## Results

To investigate the dynamics of *P. falciparum* proliferation inside human red blood cells, we developed fluorescent reporter cell lines, which permit, e.g., the tracking of DNA replication and nuclear division events in the schizont stage over time (Klaus *et al*., 2022). These reporter cell lines can be combined with fluorescent dyes to investigate the dynamics of additional cellular structures over time (Machado *et al*., 2023). Yet, when we paired fluorescent reporter lines with the lipid dye BODIPY TR Ceramide (Marks *et al*., 2008), we noticed a marked susceptibility of the cells to illumination during imaging. To quantify this effect, we compared RBC survival during imaging with and without BODIPY TR Ceramide staining (Fig. 1).

**Fig. 1:**
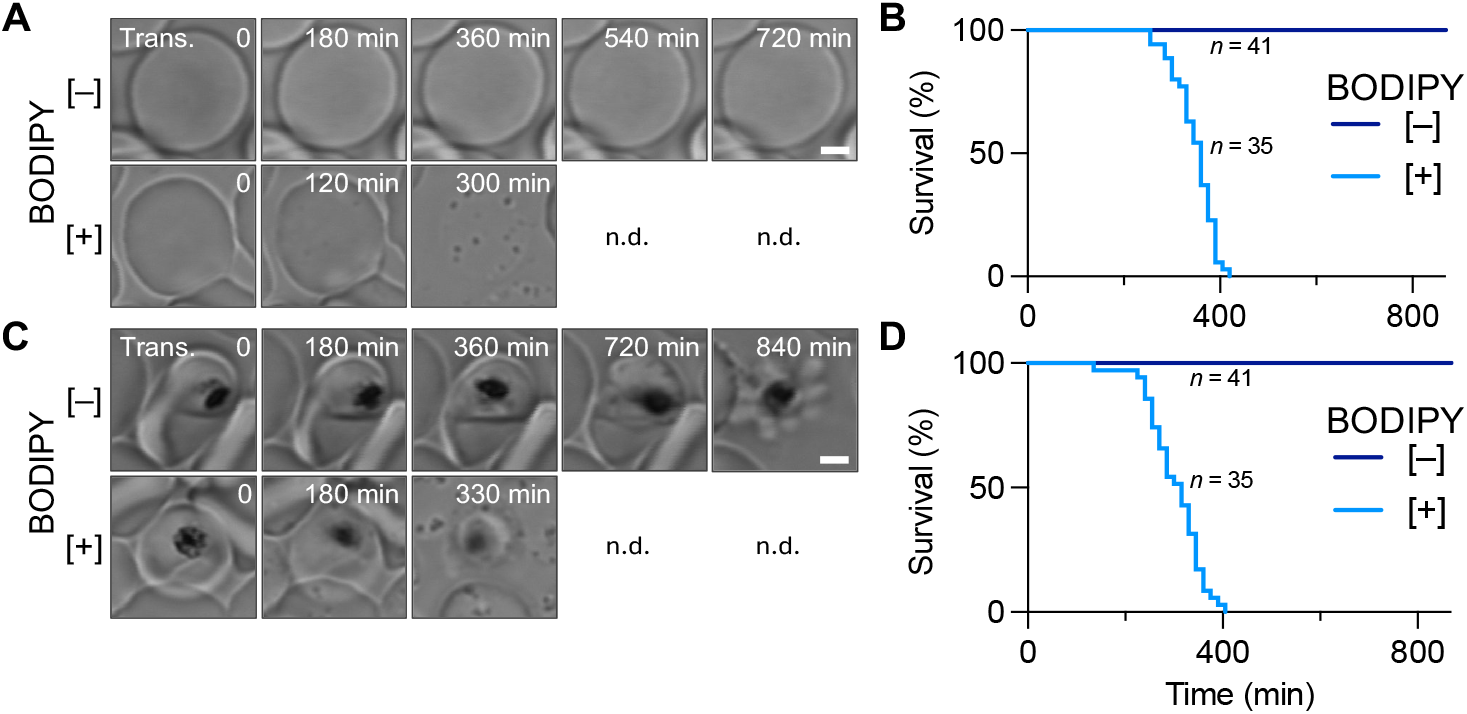
BODIPY TR Ceramide staining affects red blood cell survival during live-cell imaging. **A** Time-lapse microscopy of red blood cells, unstained [–] and stained [+] with BODIPY TR Ceramide. **B** Survival of red blood cells during imaging. **C** Time-lapse microscopy of red blood cells infected with *P. falciparum*, unstained [–] and stained [+] with BODIPY TR Ceramide. **D** Survival of red blood cells infected with *P. falciparum* during imaging. Scale bars, 2 μm.

Using relatively gentle imaging conditions (i.e., 0.2% laser power for BODIPY TR Ceramide and 0.1% for transmission image detection, z-stack with 11 slices every 15 min), we found that none of the unstained cells demised, regardless of whether they were *P. falciparum*-infected or not. In contrast, BODIPY TR Ceramide-stained cells were all lysed after approx. 400 min of imaging, with a slightly earlier onset of lysis in infected cells. Increased laser power during imaging led to a much earlier RBC lysis in both uninfected and infected cells (Fig. S1), supporting previous reports of phototoxicity associated with BODIPY-based dyes (Kähärä *et al*., 2022).

This observation in mind, we serendipitously noticed that a 32-amino acid peptide derived from *P. falciparum* PCNA1 (33 amino acids including the methionine from the start codon) appeared to target GFP to membranes in fixed cells (Fig. 2A). Previously, we have shown that PCNA1 transiently accumulates in the subset of nuclei of a multinucleated schizont that are replicating their DNA (Klaus *et al*., 2022; Machado *et al*., 2023). To investigate its nucleocytoplasmic transport, we fused multiple predicted Nuclear Localization Signals (NLS) and Nuclear Export Signals (NES) of PCNA1 to GFP via a flexible (GGGGS)2 linker (Chen *et al*., 2013). These fusion proteins were episomally expressed under the control of the *P. falciparum* CRT promoter and the *P. berghei* DHFR/TS terminator. Assessing their sub-cellular localization, we found that the predicted NLS1::GFP fusion did not localize to the nucleus, but appeared to localize to membranes (Fig. 2A, B). Given the phototoxic effects of BODIPY TR Ceramide staining, especially during long-term live-cell imaging (Fig. 1, Fig. S1), we assessed whether this 32-amino acid peptide can serve as an alternative **F**ixed and **L**ive-cell **U**niversal **M**embrane **M**arker for **I**maging (FLUMMI) (Fig. 2A, B).

**Fig. 2:**
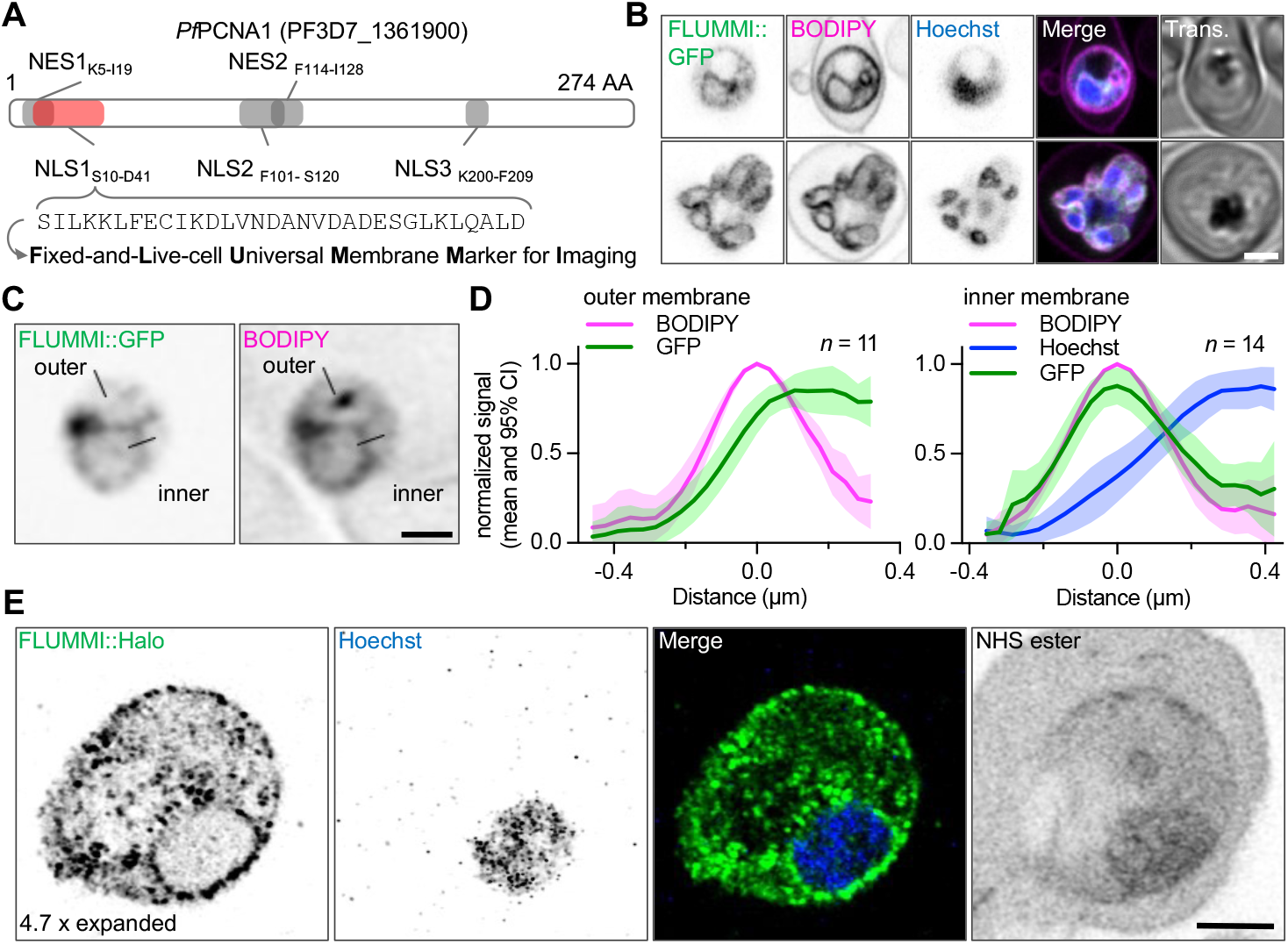
A 32-amino acid peptide targets proteins to internal membranes in fixed cells. **A** Illustration of the position of predicted nuclear localization (NLS) and nuclear export signals (NES) in the sequence of PfPCNA1. The primary sequence of the 32-amino acid long predicted NLS1 is shown below, which constitutes the Fixed– and Live–cell Universal Membrane Marker for Imaging, FLUMMI. **B** In fixed cells, fusion of FLUMMI to GFP showed a signal proximal to membranes, similar to the membrane marker BODIPY TR Ceramide; scale bar, 2 μm. **C** Comparison of FLUMMI::GFP and BODIPY TR Ceramide signal intensities. Lines highlight representative positions of line plots for outer and inner membranes; scale bar, 1 μm. **D** Quantification of FLUMMI::GFP and BODIPY TR Ceramide signal intensities (min-max normalized, with the maximal BODIPY TR Ceramide value set at distance 0 μm). For inner membranes, the signal of the DNA dye Hoechst 33342 was also plotted. Line, mean; band, 95% CI. **E** Ultrastructure expansion microscopy (U-ExM) of a cell line expressing FLUMMI fused to a Halo-tag and detected by an anti-Halo antibody; scale bar, 4 μm.

Structure prediction of FLUMMI::GFP employing AlphaFold 3 (Abramson *et al*., 2024) indicated a short α-helical structure followed by a disordered region, which stretches over the linker peptide into the structured GFP (Fig. S2A). We noted that the FLUMMI alpha helix was longitudinally split into a hydrophilic and hydrophobic side (Fig. S1B), potentially resulting in an amphiphilic peptide. We also tested if FLUMMI resembles a signal peptide, which would target the fusion protein to the ER and the secretory pathway. Using the online peptide prediction tool SignalP6 (Teufel *et al*., 2022), we found that the likelihood of FLUMMI being a functional signal peptide was with < 0.001% very low, suggesting that FLUMMI does not localize to membranes via the ER. In addition, the apparent membrane targeting was not observed when expressing GFP alone (Fig. S2 C).

Next, we quantified the signal intensities of FLUMMI::GFP and BODIPY TR Ceramide in chemically fixed *P. falciparum* cells (Fig. 2C). While BODIPY TR Ceramide was superior in marking the outer membranes (likely the plasma membrane and parasitophorous vacuole membrane), we found that FLUMMI::GFP marked the nuclear envelope equally well (Fig. 2D).

To test if FLUMMI permits a marking of membranes in ultrastructure expansion microscopy (U-ExM) (Gambarotto *et al*., 2019; Bertiaux *et al*., 2021; Liffner and Absalon, 2021), we fused it to a Halo tag, which we detected with an antibody. In expanded cells, FLUMMI::Halo showed a similar subcellular localization like FLUMMI::GFP in unexpanded cells (Fig. 2E). Previously, BODIPY TR Ceramide has been successfully used in U-ExM (Liffner and Absalon, 2021; Liffner *et al*., 2023; Liffner and Absalon, 2024).

In our hands, however, it showed an inferior signal compared to FLUMMI::Halo (Fig. S3), potentially due to signal amplification by antibody staining.

Subsequently, we assessed FLUMMI::GFP during live-cell imaging. Similar as in fixed cells (Fig. 2B–E), short-term live-cell imaging revealed that FLUMMI::GFP localized predominantly to internal membranes, comparable to BODIPY TR Ceramide staining (Fig. 3A). Notably, during long-term live-cell imaging, FLUMMI::GFP expressing parasites did not lyse, and although the GFP intensity varied from cell to cell, it remained relatively stable over 800 min of imaging (Fig. 3B, C). Since FLUMMI::GFP was expressed from an episomal plasmid, we also tested if selection for cells with a higher plasmid copy number translated into an increased FLUMMI::GFP signal (Epp *et al*., 2008). Thus, we increased the drug selection pressure by culturing FLUMMI::GFP expressing parasites in the 10-fold WR99210 concentration (i.e. 25 nM). On average, this led to an approximately 25-fold brighter GFP fluorescence (Fig. 3C), suggesting that signal intensity can be adapted to experimental needs by modulating the drug selection pressure.

**Fig. 3:**
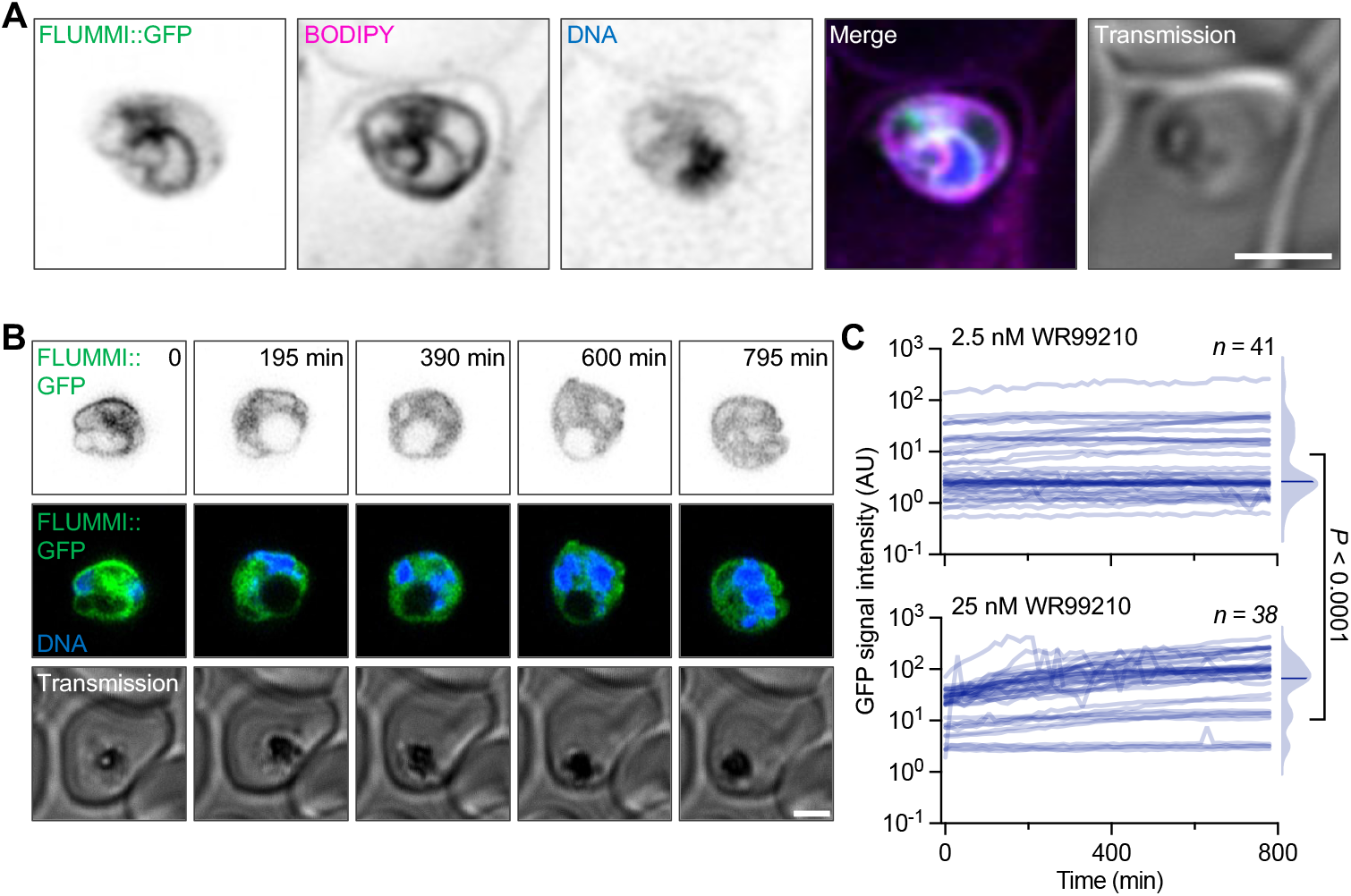
FLUMMI::GFP marks membranes in short-term and long-term live-cell imaging. **A** Short-term live-cell imaging FLUMMI::GFP expressing parasites co-stained with BODIPY TR Ceramide and the DNA dye SPY650-DNA. **B** Long-term live-cell imaging of FLUMMI::GFP expressing parasites co-stained with the DNA dye SPY650-DNA. **C** Quantification of the FLUMMI::GFP signal intensity over time. Increasing the drug pressure for the presence of the FLUMMI::GFP expressing and WR99210-resistance conferring plasmid selected for on average approx. 25–fold brighter parasites. Half violins, average values of individual parasites over time; bar, median; *P-*value, Mann Whitney test. Scale bars, 2 μm.

Together, this suggests that FLUMMI could be a substitute for BODIPY TR Ceramide staining to mark internal membranes during long-term live-cell imaging of *P. falciparum* blood stages.

Next, we tested if FLUMMI could also serve as a membrane marker in other *Plasmodium* species, as well as other life cycle stages. In the rodent malaria parasite *P. berghei*, we integrated a FLUMMI::mScarlet construct in the silent locus on chromosome 6 (SIL6) (Kooij *et al*., 2012) (Fig. S4A, B). Expression of FLUMMI::mScarlet from this endogenous locus and driven by the *hsp70* promoter led to a slight but significant reduction in parasite growth rate from an 8-fold to a 6-fold increase over 24 hours (Fig. 4A, Fig. S4). Similar to *P. falciparum*, FLUMMI::mScarlet localized to internal membranes, marked with BODIPY FL, in both fixed and live *P. berghei* asexual blood stages (Fig. 4B).

**Fig. 4:**
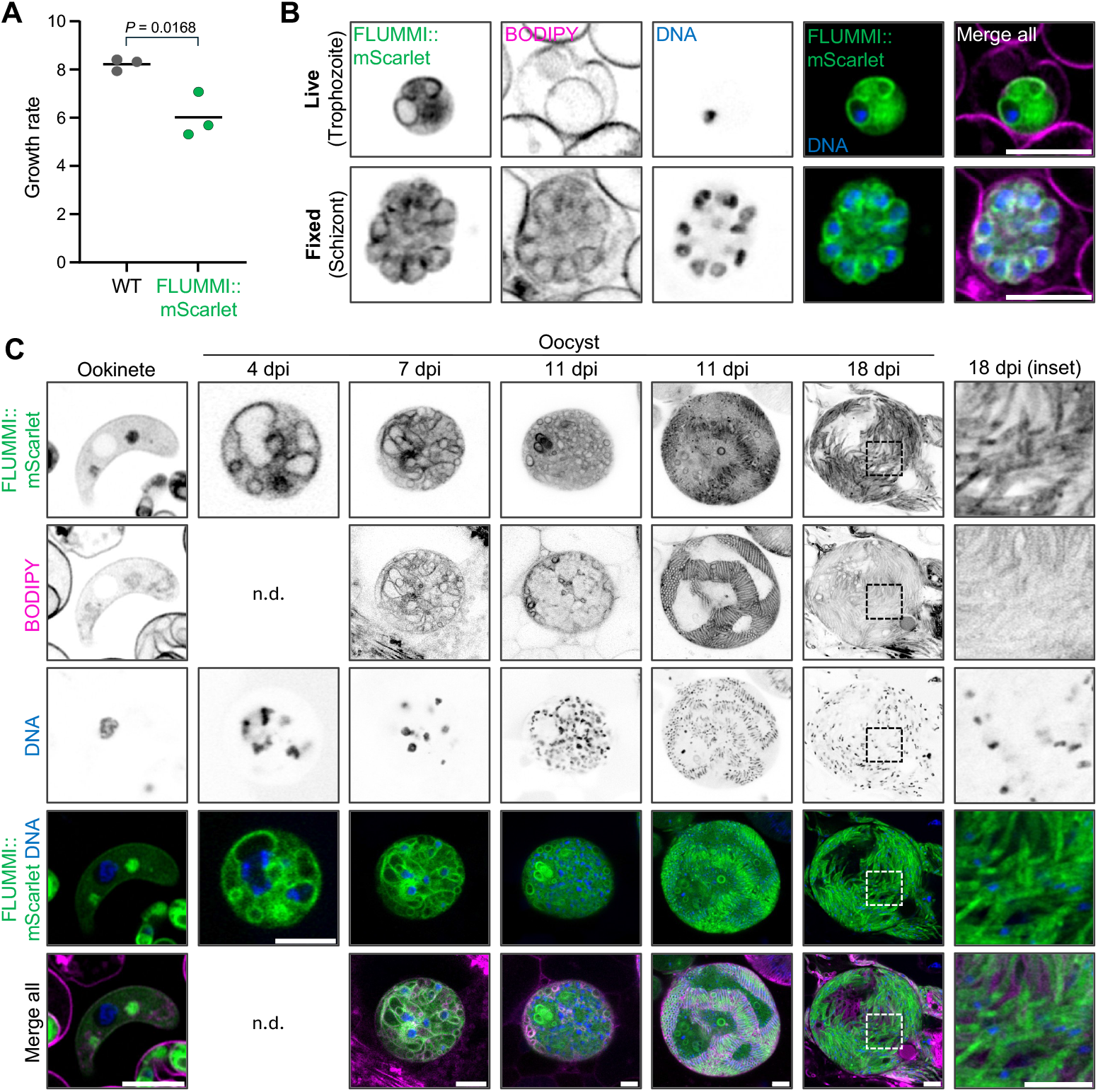
FLUMMI marks membranes throughout *Plasmodium berghei* development in the vertebrate and the *Anopheles* mosquito. **A** *P. berghei* parasites expressing FLUMMI::mScarlet endogenously from the SIL6 locus show a slightly reduced growth in asexual blood stages. Bar, mean; *P*-value, unpaired two-tailed t-test. **B** Fixed- and live-cell short-term imaging of FLUMMI::mScarlet-expressing asexual blood stage *P. berghei* parasites, co-stained with BODIPY FL and the DNA dye Hoechst 33342. **C** Live-cell imaging of different mosquito-stage *P. berghei*, expressing FLUMMI::mScarlet and co-stained with BODIPY FL and the DNA dye Hoechst 33342. Scale bars, 5 μm.

We then followed FLUMMI localization in mosquito-infecting stages of *P. berghei*. In ookinetes, FLUMMI::mScarlet appeared to localize predominantly to two distinct spots, possibly the crystalloids, which were also detectable with BODIPY FL (Fig. 4C). The crystalloids are specific organelles rich in membranes and membrane-associated proteins such as a palmitoyl-*S*-acyl transferase or the multipass transmembrane protein NAD(P) transhydrogenase (Saeed *et al*., 2015; Santos *et al*., 2016; Saeed *et al*., 2020). Notably, BODIPY FL also did stain ookinetes weaker than asexual blood stages. In early oocysts (day 4 post infection), we observed an extended internal network of membranes, including the nuclear membrane and possibly the ER (Fig. 4C). We noted that the nuclear compartment, as identified by FLUMMI::mScarlet, appeared often much larger than the signal derived from the DNA dye, indicating a level of intranuclear compartmentalization in these polyploid nuclei. At later stages of oocyst development, some of the internal membrane structures seemed not to be stained with BODIPY FL, or only weakly, possibly because of difficulties of the dye to penetrate the oocyst capsule, highlighting the advantage of using a genetically encoded membrane marker over an externally added dye. In sporulating oocysts, FLUMMI::mScarlet additionally localized to foci adjacent to the nucleus of the sporozoites, potentially marking the nucleus associated ER (Fig. 4C).

Finally, we aimed to test if FLUMMI was able to mark membranes in cells from evolutionary distant organisms. To this end, we transfected human HeLa-derived TZM-bl cells (Wei *et al*., 2002) with a FLUMMI::GFP expression construct. Fluorescence microscopy revealed a homogenously distributed signal in the cells’ cytoplasm and nucleus (Fig. S5A-C). This distribution was reminiscent to that of GFP alone and Western Blot analysis confirmed that the majority of ectopically expressed protein in these cells was GFP without the 32 amino acid PCNA1-derived peptide. The lack of detection of full length FLUMMI::GFP in these cells may reflect, e.g., protein degradation and/or the use of an alternative start codon. While more human cell types need to be tested for a generalized conclusion, this result suggests that expression of FLUMMI::GFP is subject to different regulation in *Plasmodium* spp. versus human cells and, thus, may serve as a *Plasmodium*-specific membrane marker.

Together, our data indicate that FLUMMI can be used to track the dynamic development of internal membranes in different developmental stages of *Plasmodium*.

## Discussion

We here report the serendipitous discovery of a genetically encoded membrane marker, FLUMMI, which allows visualization of internal membranes both in fixed and in live *Plasmodium* cells. Importantly, expression of FLUMMI::GFP permits long-term live cell imaging of membrane dynamics over at least 12 h, avoiding the cytotoxicity observed with BODIPY TR Ceramide under the same conditions. Although not readily usable like a dye, FLUMMI marks only membranes of the parasite without contributing signal from the host cell. While the membrane localization can be attributed to the FLUMMI peptide (rather than the GFP (Fig. S2) or the frequently used (GGGGS)x linker (Chen *et al*., 2013; Ambekar *et al*., 2022; Klaus *et al*., 2022)), the mechanism by which FLUMMI targets protein tags to membranes is unknown and remains an interesting question for future research.

Modeling of its structure revealed a possibly amphiphilic alpha helix that might bind laterally to the membrane. However, the non-uniform staining of *Plasmodium* membranes (Fig. 2D) suggests a dependence on specific factors. These may be the lipid composition or specific membrane-associated proteins. Albeit unlikely, FLUMMI may also resemble a non-canonical signal peptide, which would target proteins to the ER and secretory pathway. Thus, for fixed and short-term live microscopy, FLUMMI and BODIPY might complement each other, highlighting different aspects of parasite membrane composition and architecture. Especially in *Plasmodium* oocysts – parasite stages that, due to their thick proteinaceous cyst wall, are notoriously hard to stain with external dyes – FLUMMI revealed a previously underappreciated extensive network of internal membranes. This network encompasses likely nuclear membranes and the ER, but possibly includes other organelles, too.

Notably, FLUMMI expression in oocysts indicates that the DNA occupies only a small fraction of the entire nuclear volume. For the polyploid nuclei of an oocyst, an intranuclear compartmentalization may be a means to organize its multiple genomes. This exemplifies how this membrane reporter could assist in elucidating the little-understood cell biology of early oocyst development.

In conclusion, we here show that FLUMMI can serve as a membrane marker for fluorescence imaging in the malaria-causing parasite *Plasmodium*, in particular for long-term live cell microscopy. FLUMMI is also compatible with advanced imaging techniques like ultrastructure expansion microscopy and works across *Plasmodium* species and parasite stages, rendering it a new tool to study membrane dynamics.

## Experimental Procedures

### *P. falciparum* cell culture

*P. falciparum* 3D7 is a laboratory-adapted strain, derived from the isolate NF54, which originated from a case of airport malaria in the Netherlands (Ponnudurai *et al*., 1981). *P. falciparum* 3D7 was cultivated as previously described (Trager and Jensen, 1976). In brief, we used fresh O, Rh+ erythrocytes at 2−4% hematocrit in supplemented RPMI 1640 medium (with 0.2 mM hypoxanthine (CCPro), 25 mM HEPES, pH 7.4 (Merck), and 12.5 μg/ml gentamicin (Carl Roth), as well as 0.5% AlbuMAX II (gibco)), at 37 °C in 90% relative humidity, 5% O2, and 3% CO2. Parasitemia was monitored via blood smears that were fixed with 100% methanol for 10 seconds and stained for 1 s in Hemacolor solution II (VWR) and subsequently for 20s in Hemacolor solution III (VWR) to visualize red blood cells and parasites using either a Nikon Eclipse E100 or a Zeiss Axiostar Plus microscope with a 100× oil immersion objective. Routine synchronizations were performed by 5% sorbitol treatment as previously described (Lambros and Vanderberg, 1979).

### Cloning of plasmids for *Plasmodium* transfection

To investigate the 32-amino acid predicted PfPCNA1 NLS1/FLUMMI peptide in *P. falciparum*, we expressed FLUMMI with an additional start codon and fused to eGFP using a PP(GGGGS)2 linker (with PP deriving from an AvrII site used for molecular cloning) from the episomal plasmid pARL under the control of the *P. falciparum* CRT promoter and the *P. berghei* DHFR/TS terminator (pARL_NLS1::GFP). Additionally, this plasmid contained a WR99210 resistance cassette (Fidock and Wellems, 1997). The resulting parasite line was used for fixed and live cell imaging. For Ultrastructure Expansion Microscopy (U-ExM), we fused FLUMMI to a HaloTag in the pARL plasmid backbone. All plasmids were sequenced by Sanger sequencing before transfection.

To express FLUMMI in *P. berghei*, the plasmid pBAT-SIL6-MCS (Kooij *et al*., 2012), was digested with BamHI and NdeI. The putative NLS sequence of *P. falciparum* PCNA1 was amplified from pARL_NLS1::GFP and mScarlet was amplified from the Addgene Plasmid 85054. Fragments were combined using Gibson assembly using the NEB HiFi Assembly kit (NEB). All plasmids were sequenced by Sanger sequencing before transfection. All primers and plasmids used to generate these cell lines are available upon request.

### Transfection of *P*. *falciparum*

Transgenic parasites were generated as previously described (Rug and Maier, 2012). In brief, 75–100 μg purified plasmid DNA was resuspended in 30 μl of TE buffer [10 mM tris (pH 8.0) and 1 mM EDTA (pH 8.0)] and added to 370 μl of CytoMix [120 mM KCl, 0.15 M CaCl2, 2 mM EGTA, 5 mM MgCl2, 10 mM K2HPO4/KH2PO4, and 25 mM Hepes (pH 7.6)].

The DNA/CytoMix solution was added to 200 μl packed erythrocytes infected with ring-stage *P. falciparum* 3D7 parasites at a parasitemia of 5–8%. Electroporation was performed in a 0.2 cm gap electroporation cuvette (Bio-Rad) with a Gene Pulser II (Bio-Rad, settings: 0.31 kV, 0.950 μF, capacitance set to “High Cap,” resistance on the Pulse Controller II set to “Infinite”). Immediately after electroporation the sample was added to a 10 ml petri dish and cultured under standard conditions. To select for transgenic parasites, we used 2.5 nM WR99210 (kind gift of Jacobus Pharmaceutical Company, Princeton, USA).

### *P. falciparum* preparation for fixed-cell imaging

Fluorescence staining for microscopy of chemically fixed blood-stage parasites was done as previously described with minor alterations (Simon *et al*., 2021; Machado *et al*., 2023). In brief, we seeded cells on ibidi Treat 8-well dishes, rinsed them with pre-warmed PBS and subsequently fixed with pre-warmed 4% PFA/PBS for 20 minutes at 37 °C. After fixation, cells were washed with PBS and permeabilized with 0.1% Triton X-100/PBS for 15 minutes at room temperature (RT). Cells were then washed two times with PBS, leaving 200 μl of PBS in each well for imaging. Additionally, Hoechst 33342 was added approximately 1 hour before imaging to label the nuclei, while BODIPY TR Ceramide (Thermo Fisher Scientific, D7540) was added 20 minutes prior to imaging to visualize membranes. Endogenous GFP signal was strong enough to not require additional antibody staining before imaging.

### *P. falciparum* preparation for live-cell imaging

For live-cell imaging, infected red blood cells were seeded in either a round 35 mm imaging dish or an ibidi 8-well dish. When using ibidiTreat-coated dishes, the dishes were rinsed twice with PBS before seeding. Uncoated dishes were coated prior to seeding with Concanavalin A (200 μl for round dish; 80 μl/well for an 8-well dish) and incubated at 37 °C for 20-30 minutes. Excess of concanavalin A was removed, and dishes were rinsed with PBS twice.

To prepare the cells, 500 μl of resuspended cell culture was pelleted (800 g for 2 min) and washed twice with supplemented RPMI 1640 without AlbuMAX II. Cells were resuspended in 1 ml of pre-warmed supplemented RPMI 1640 without AlbuMAX II, and 125 μl per 8-well or 250 μl per 35 mm round dish was used for seeding. To allow for cell attachment, dishes were incubated for 10 minutes at 37 °C. To create a monolayer, the cells were gently washed with and kept under normal culture conditions in supplemented RPMI 1640 without AlbuMAX II until the medium was changed to pre-warmed imaging medium (i.e., supplemented RPMI 1640 without phenol red (PAN-Biotech). The SPY650-DNA dye was added to the imaging medium 2 hours prior to imaging and BODIPY TR Ceramide was added 20 min prior to imaging.

### *P. falciparum* preparation Ultrastructure-Expansion Microscopy (U-ExM)

Ultrastructure Expansion Microscopy was performed as previously described (Gambarotto *et al*., 2019; Bertiaux *et al*., 2021; Liffner and Absalon, 2021). In brief, 12 mm coverslips were coated with poly-D-Lysin for 1 hour at 37 °C and then washed three times with PBS. PBS was removed and coverslips were stored at 4 °C in a humid chamber. *P. falciparum*-infected RBCs were enriched by magnetic affinity purification using a MACS LS column (Miltenyi Biotec), the eluate was pelleted (800 g for 2 min) and the supernatant was removed. Cells were resuspended in 300 μl supplemented RPMI 1640 without AlbuMAX II and 100 μl of the suspension were added to the coverslip. After the seeding for 10 minutes at 37 °C, the supernatant was removed and the infected RBCs were chemically fixed by adding 50 μl of 4% PFA/PBS followed by 20 min incubation at 37 °C. Then, coverslips were washed with PBS and stored in a 12-well plate in PBS. To later permit protein anchoring to the poly acrylamide gel, we added 1 ml formaldehyde acrylamide (FA/AA) solution to the cells and incubated at 37 °C for 3.5 to 5 hours with gentle agitation on an orbital shaker plate at 80 rpm.

Gelation was carried out on ice in a humid chamber by dipping the coverslips in PBS and then placing them on a 35 μl drop of monomer solution with tetramethylethylendiamine (TMED, Thermo Fisher Scientific) and ammonium persulfate (APS, Thermo Fisher Scientific) with cells facing the monomer solution mixture. After incubation for 5 minutes on ice, the humid chamber was covered with aluminum foil and incubated for 1 hour at 37

°C. Coverslips were transferred with the gel facing up in a 6-well plate containing 2 ml denaturation buffer and were incubated until the gel detached from the coverslip. Then, gels were transferred into a 1.5 ml reaction tube with fresh denaturation buffer and kept for 1.5 hours at 95 °C. Denaturation buffer was removed, and gels were left for the first expansion in ddH2O. After 30 minutes, the water was replaced by fresh water supplemented with sodium acetate and left overnight at RT. Next, the gel size was measured to determine the expansion factor. For immunofluorescence staining, the gels were cut and incubated in PBS twice for 15 minutes, which leads to shrinkage of the gels. PBS was replaced by 3% BSA/PBS and incubated for 30 minutes at RT, then the primary antibody was added (250 μl, 1:1000 Anti-halo, Promega) and incubated for three days at 4 °C with gentle agitation on an orbital shaker plate. Next, the gels were washed five times for 10 min with 0.5% Tween/PBS and the secondary antibody solution (anti-mouse Alexa Fluor 488, 1:500, Invitrogen) with Hoechst 33342 (1:100 Thermo Fischer Scientific) was added and incubated for 2.5 hours in the dark with gentle agitation at 37 °C. After five washes with 0.5% Tween/PBS for 10 min each, Atto 647 NHS ester (Atto-Tech) staining was performed for 1.5 hours at RT with gentle agitation. After washing as described above, the gels were placed in water supplemented with BODIPY TR Ceramide (2 μM) for the second expansion and for membrane staining. For imaging, Ibidi Glass bottom dishes were coated with poly-D-Lysine and gels were placed in the wells with cells facing downwards.

### Expression and Localization of FLUMMI::GFP in mammalian cells

HeLa-derived TZM-bl cells (Wei *et al*., 2002) were transfected with pN1-GFP or pFLUMMI::GFP expression constructs using Lipofectamine 2000 (Thermo Fisher Scientific) according to manufacturer’s protocol. In brief, 5 × 104 TZM-bl cells were seeded on glass coverslips overnight and then transfected using 1 μg of DNA and 2 μl of lipofectamine. Samples were then harvested 48h post transfection, the cells were washed in 1x PBS, and the plasma membrane was stained using WGA-647 (Invitrogen) at a final concentration of 1 μg/ml for 5 min at RT. The samples were then fixed using 3% PFA for 15 min in the dark and further processed for light microscopy analysis. Point laser scanning confocal microscopy was then performed on a Leica SP8 microscope using an HC PL APO CS2 63×/1.4 N.A. oil immersion objective. Images were acquired using PMT detectors in reflection mode with a laser excitation at 488 or 633 nm, and spectral detection windows set between 500 nm to 570 nm, or 650 to 700 nm wavelengths.

Western blot analysis was done according to standard procedures. In brief, TZM-bl were lysed with 2× SDS sample buffer (10% glycerol, 6% SDS, 130 mM Tris HCl pH 6.8, 10% β-mercaptoethanol) supplemented with protease/phosphatase inhibitor (Thermo Fisher Scientific, 78440) and boiled for 5 min at 95 °C. Protein samples were resolved by SDS-PAGE electrophoresis and blotted onto nitrocellulose membranes. Membranes were blocked in 5% milk/PBST for 1 h and were incubated with primary antibodies overnight at 4 °C (rat anti-GFP: 1:1000, Chromotek, 3H9; mouse anti-GAPDH: 1:5000, Biorad, MCA4740). After three washes with 5% milk/PBST, membranes were incubated with HRP-conjugated secondary antibodies (Goat anti-mouse IgG-HRP: 1:5000, 31431; rabbit anti-rat IgG-HRP: 1:5000, PA1-28786; Invitrogen) for 1 h at room temperature. Membranes were again washed three times and visualized by chemiluminescence.

### Ethics statement and mice

All experiments were performed in accordance with GV-SOLAS and FELASA guidelines and have been approved by the German authorities (Regierungspräsidium Karlsruhe). Female Swiss mice weighing 20-25 g were purchased from JANVIER and were aged five to ten weeks at the time point of infection. For each experiment, mice were age-matched and were allocated randomly to each group. Mice were kept in groups of 2 to 4 mice per cage under specified pathogen-free conditions within the animal facility at Heidelberg University on a 12-hour light/dark cycle at 22 °C (± 2 °C) and 50-60% humidity with *ad libitum* access to food and water.

### General maintenance of parasites and mosquitoes

Mice were typically infected with *P. berghei* by intraperitoneal injection of cryostabilates, consisting of 100 μl parasitic blood and 200 μl freezing solution (Alsever solution plus 10% glycerol), or by intraperitoneal or intravenous fresh blood transfer of a defined number of parasites. Parasitemia was determined using Giemsa-stained blood smears. Parasitized blood was collected from infected mice by cardiac puncture of fully anesthetized mice (after intraperitoneal application of ketamine/xylazine). This was also the route to euthanize the mice. Mosquitoes (*Anopheles stephensi*) were reared and maintained according to established protocols.

### Generation of transgenic *P. berghei* parasites

Transgenic *P. berghei* parasite lines were generated largely as previously described (Janse *et al*., 2006; Schwach *et al*., 2015). Briefly, 10 μg plasmid DNA was digested over night with PvuI, precipitated using ethanol and resuspended in 10 μl PBS. Schizonts were obtained by culturing 500 μl blood containing > 1% parasitemia in schizont medium (RPMI-1640, gibco) supplemented with 20% FCS (gibco) and 1 μg/ml gentamycin (PAA) for about 20 h at 37 °C and were purified over a 55% Nycodenz (Axis-shield diagnostics) gradient. DNA and schizonts were mixed with 100 μl Nucleofector solution from the human T cell Nucleofector Kit, electroporated using the Amaxa Nucleofector II device (Lonza), and immediately injected intravenously into a mouse. Transgenic parasites were selected by administering pyrimethamine (7 μg/ml, Sigma-Aldrich) to the drinking water one day after transfection. Blood-stage positive mice were bled by cardiac puncture and blood was used for preparation of cryostabilates and for genotyping. To this end, blood containing at least 1% parasitemia was lysed using 0.093% saponin and the parasite pellet was resuspended in 200 μl PBS. gDNA was isolated using the DNeasy Blood & Tissue Kit (Qiagen) according to the manufacturer’s instructions. Parasites were genotyped by PCR for amplification of the wildtype locus (locus), 5’ integration (5’int), and 3’ integration (3’int) sites.

### Determination of *P. berghei* asexual growth rate

To determine parasite growth, three mice per line were infected intravenously with 1000 iRBC. Parasitemia was assessed from day 4 to day 7 post infection by Giemsa-stained blood smears. For each mouse, the parasite growth rate was determined from the slope of a linear regression fitted to the log-transformed parasitemia curve (excluding values for day 7, where parasitemia plateaued).

### Live and fixed-cell imaging of asexual *P. berghei* parasites

To perform life-cell imaging, a drop of *P. berghei* parasitized mouse blood collected from the tail vein was stained with BODIPY FL C5-Ceramide (Thermo Fischer Scientific, D3521, 1:1000 dilution) for 1 h at 37 °C and with Hoechst-33342 (final concentration 10 μg/ml) for 5 min at 37 °C, washed once and imaged on an inverted Zeiss LSM900 microscope equipped with an Airyscan 2 detector (see below).

To fix cells for imaging, 50 μl of blood was mixed with 200 μl of RPMI medium, and immediately fixed by adding an equal volume of 4% PFA/0.0075% glutaraldehyde in PBS. Cells were incubated for 30 min at 37 °C, washed once in PBS and stored in 3% BSA/PBS at 4 °C. Prior to imaging, cells were stained with Bodipy FL (1:1000 dilution) for 1 h at RT and with Hoechst 33342 (final concentration 10 μg/ml) for 5 min at RT, washed once and imaged.

### Ookinete culture

To obtain ookinetes for live-cell imaging, mice were infected with 2 × 107 iRBC i.p.. Three days later, mice were bled, and the gametocyte-containing blood transferred to 10 ml ookinete medium (RPMI supplemented with 20% (v/v) FCS, 50 μg/ml hypoxanthine, and 100 μM xanthurenic acid, pH 7.8 – 8.0). Parasites were cultured at 19 °C for 20 to 24 hours, stained for 1h with Bodipy FL (1:1000 dilution) and for 5 min with Hoechst-33342 (final concentration 10 μg/ml) at 19 °C, washed once in ookinete medium and immediately imaged.

### Mosquito infections

Mosquitoes were infected by blood meal on a mouse three days post infection with 2 × 107 iRBC i.p.. To this end, the mouse was anesthetized by administering ketamine/xylazine solution (20 mg/ml ketamine, 0.6 mg/ml xylazine in PBS) i.p. at 5 μl/g body weight and placed onto a mosquito cage containing approximately 150-250 mosquitoes. Mosquitoes were allowed to feed for approximately 30 min, with a change of mouse position after 15 min. After feeding, mosquitoes were immediately kept at 21 °C and 80% humidity, and fed with 10% (v/v) saccharose with 0.05% PABA and 1% (v/v) NaCl.

At selected time points, mosquito midguts were dissected into PBS, stained with Bodipy FL (1:1000 dilution) and Hoechst 33342 (final concentration 10 μg/ml) for 1 h at RT, washed once and imaged. For oocysts on day 4, only staining with Hoechst 33342 was performed.

### Imaging at a Zeiss LSM900 microscope

For all imaging, unless stated otherwise, point laser scanning confocal microscopy was performed on an inverted Zeiss LSM900 microscope equipped with an Airyscan 2 detector using a Plan-Apochromat 63× /1,4 oil immersion objective and with the Airyscan detector in SR mode.

For short-term imaging of live and fixed cells, single z-stacks were recorded to cover the entire cell, ranging from 11 to 159 confocal slices. Long-term live-cell imaging was performed at 37 °C with 5% CO2 and 5% O2, and 11 confocal slices per z-stack were recorded every 15 minutes over approximately 14 hours. Multiple positions were imaged using the “Positions” tool and the focus was stabilized using the “Definite Focus” mode. Multichannel images were acquired sequentially using the 405 nm diode for Hoechst, the 488 nm laser for GFP or BODIPY FL, the 561 nm diode for SPY555-DNA, BODIPY TR Ceramide or mScarlet, and the 640 nm laser for SPY650-DNA. The scanning was done bidirectionally and the transmitted light images were captured by the T-PMT detector. Airyscan data was processed with the ZEN Blue 3.1. software.

### Image analysis

Image analysis was performed using FIJI (Schindelin *et al*., 2012). For snapshots of a single time point (fixed cells, live-cell snapshots, and U-ExM), a single z-slice at the center of the cell was selected for analysis and visualization. Channels were converted to gray scale, and inverted to display the signal on a white background for enhanced visibility. To compare the signal intensities of BODIPY TR Ceramide and FLUMMI::GFP, we used line profiles. Lines were drawn from the extracellular space or cytosol, across the membrane, and into the cytoplasm or nucleus, respectively. Signal intensities along these lines were measured using the plot profile function of FIJI. Intensities were min-max normalized and the maximal BODIPY TR Ceramide value was set at the distance 0 μm for graphical representation. Long-term imaging data was pre-processed to correct for drift, using the 3D Drift Correction in FIJI. To quantitatively assess the GFP signal from long-term imaging data, a region of interest was first cropped out. Then the sum of the signal (i.e., the raw integrated density) of all z-slices per time point was plotted over time in arbitrary units. For Kaplan-Meier plots, long-term imaging data was used to evaluate cell survival over time. Time of cell death was determined based on loss of structural integrity, i.e., cell lysis of parasite and/or red blood cell. Further data processing and visualization were performed using Microsoft Office, Python (Jupyter notebook), and GraphPad Prism.

## Acknowledgements

The authors are grateful to Severina Klaus, Mirko Singer, and members of the Hentzschel and Ganter labs for helpful discussions; Annett Pettrich for the gift of BODIPY FL C5-Ceramide; the Infectious Diseases Imaging Platform (www.idip-heidelberg.org); and the *Plasmodium* database PlasmoDB (www.plasmodb.org), which facilitated this work. F.H. receives funding from the ERC (Starting Grant PlasmoArp, 101162759) and the German Research Foundation (Deutsche Forschungsgemeinschaft, DFG) through SPP 2225, project number 531930468. M.G. is supported by the Health + Life Science Alliance Heidelberg Mannheim, receiving state funding approved by the State Parliament of Baden-Württemberg. O.T.F. and M.G. receive support from the German Research Foundation (Deutsche Forschungsgemeinschaft, DFG) through SFB 1129, project number 240245660, subprojects TP8 and TP18, respectively.

## Author contributions

Conceptualization, L. Z., F.H. and M. G.; Methodology, L. Z., F.H. and M. G.; Investigation, L. Z., V. R., S.S.A., M.L., J.N. and A.K.; Resources, S.S.A. and K. Z.; Writing – Original Draft, L. Z., F.H. and M. G.; Writing – Review & Editing, L. Z., S.S.A., F.H. and M. G.; Visualization, F.H. and M. G.; Supervision, F.H. and M. G.; Funding Acquisition, O.T.F., F.H., and M. G.

