## Supplementary information for "A genetically encoded marker for imaging membranes of *Plasmodium*"

### Supplementary figures

#### Fig. S1

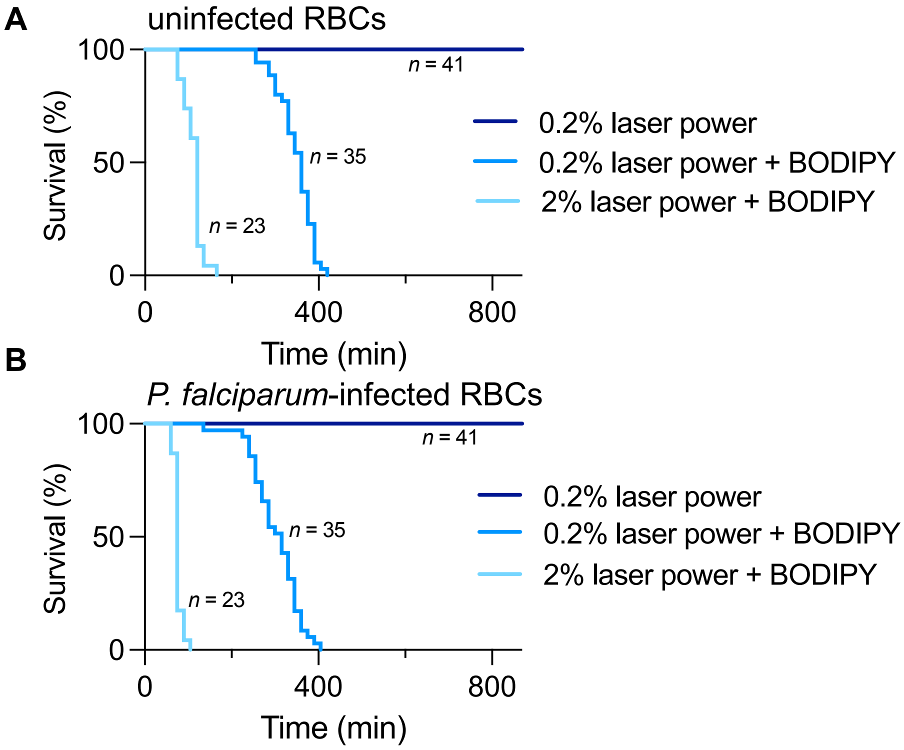


**Fig. S1: BODIPY TR Ceramide staining affects red blood cell survival during live-cell imaging.** **A** Survival of uninfected red blood cells during imaging. **B** Survival of *P. falciparum*-infected red blood cells during imaging. With 0.1% laser power for the transmission image.

### Fig. S2

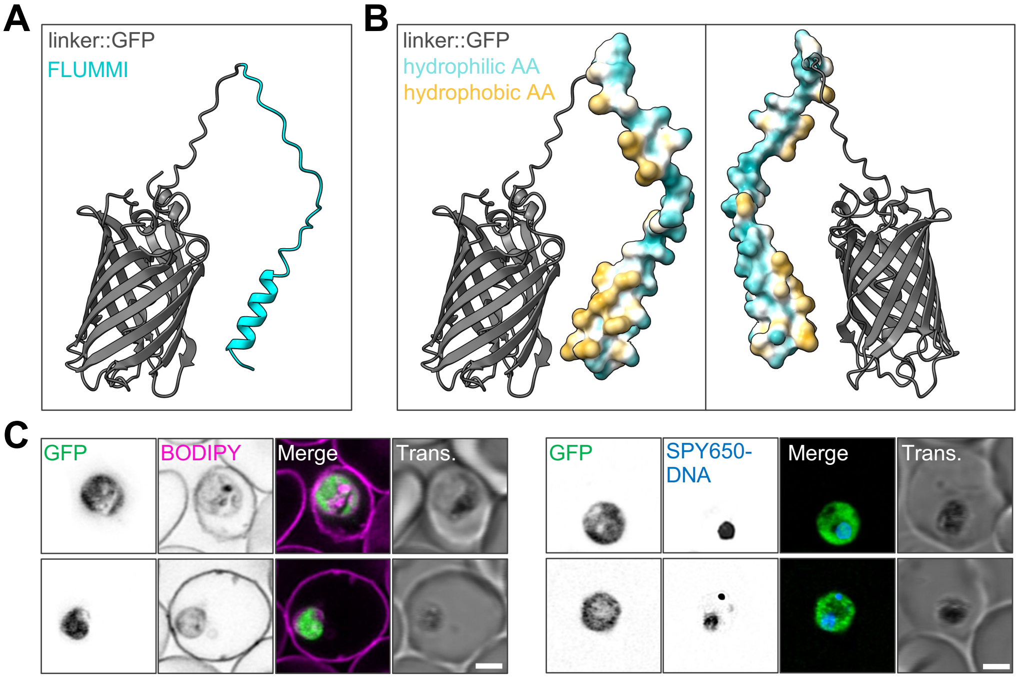


**Fig. S2: AlphaFold3 predicts FLUMMI to be an amphiphilic peptide that harbors the membrane-targeting activity. A** AlphaFold3 prediction of FLUMMI (cyan) fuse to GFP via a linker peptide (PP[GGGGS]$_2$, with PP deriving from an AvrII site for molecular cloning). **B** Surface rendering of FLUMMI with hydrophobic and hydrophilic amino acid (AA) residues indicated in yellow and cyan, respectively. **C** Short-term live-cell imaging of parasites expressing GFP co-stained with BODIPY TR Ceramide (left) or the DNA dye SPY650-DNA (right). Scale bars, 2 µm.

### Fig. S3

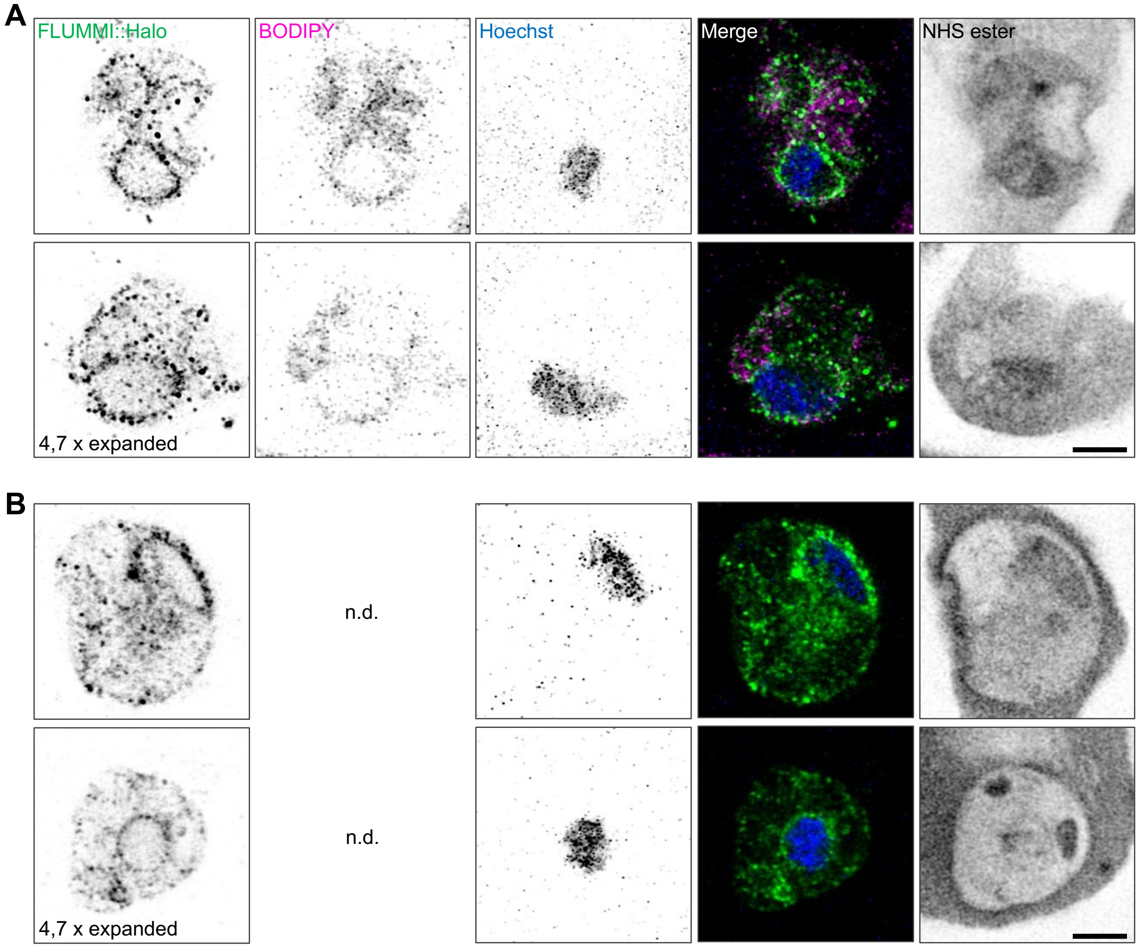


**Fig. S3: Ultrastructure expansion microscopy of FLUMMI::Halo expressing *P. falciparum*. A** FLUMMI::Halo expressing cells were co-stained with BODIPY TR Ceramide and the DNA dye Hoechst 33342 or **B** with Hoechst 33342 (bottom); scale bars, 4 µm.

### Fig. S4

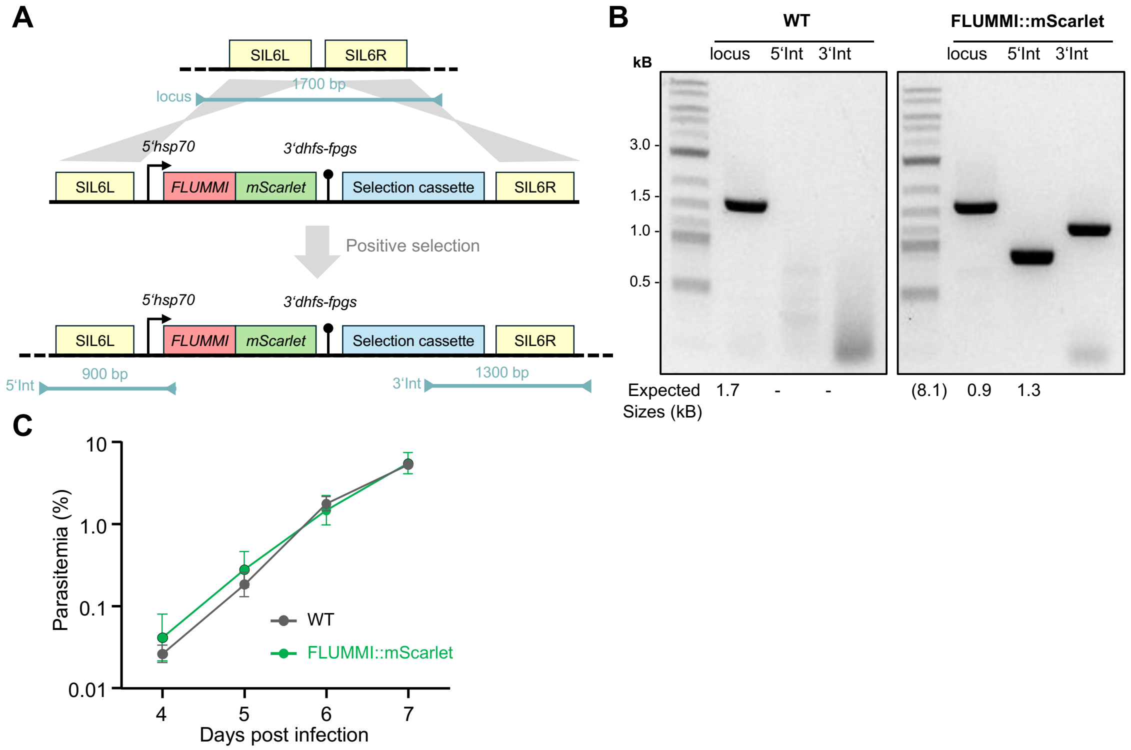


**Fig. S4: Generation and basic characterization of a FLUMMI::mScarlet expressing *P. berghei* cell line. A** Integration strategy to generate the FLUMMI::mScarlet cell line. The silent locus on chromosome 6 (SIL6) was targeted with a linearized integration construct containing two homology regions (SIL6L, SIL6R), a FLUMMI::mScarlet expression cassette, as well as a cassette conferring drug resistance (selection cassette). Upon homologous recombination, this construct is integrated in the SIL6 locus. Test primer combinations for diagnostic PCR are indicated by arrow heads and the expected fragments are shown as lines. **B** Diagnostic PCR confirms the successful integration of FLUMMI::mScarlet into the SIL6 locus. **C** Asexual blood-stage proliferation of *P. berghei* expressing FLUMMI::mScarlet was compared to wild type parasites after i.v. injection of 1000 infected erythrocytes of each cell line into recipient animals ($n$ = 3). Parasitemia was determined daily using Giemsa-stained blood smears.

**Fig. S5**

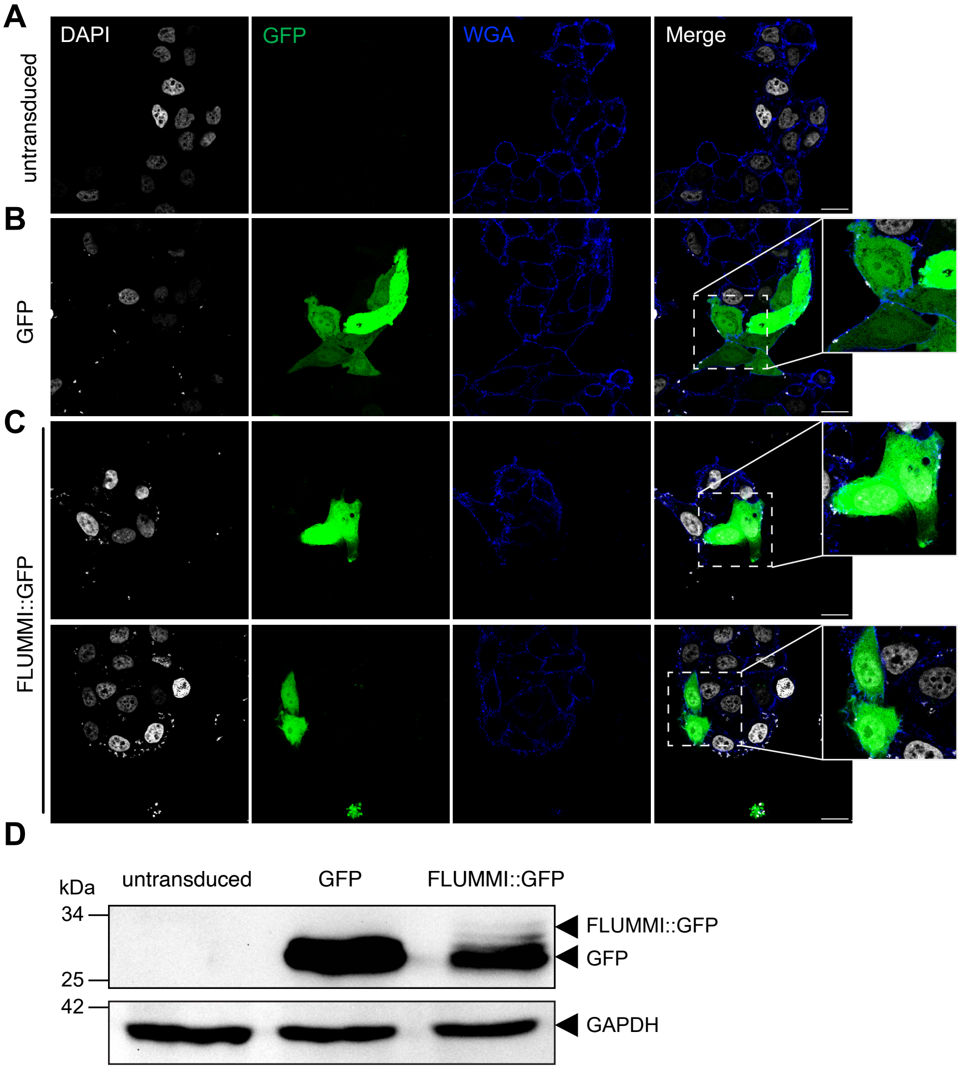


**Fig. S5: FLUMMI::eGFP shows a pancellular localization in human cells. A** Control untransduced HeLa-derived TZM-bl cells stained with DAPI to detect DNA and wheat germ agglutinin (WGA) to mark the plasmamembrane. **B** Control TZM-bl cells transduced with a GFP expressing construct and staind with DAPI and WGA; inset, highlighting the pancellular localization of GFP. **C** TZM-bl cells transduced with a FLUMMI::GFP expressing construct and staind with DAPI and WGA; inset, highlighting the FLUMMI::GFP signal akin to the control GFP signal in B. **D** Western Blot analysis of TZM-bl cells (untransduced, GFP expressing, or FLUMMI::GFP expressing), with GAPDH serving as a loading control. Scale bars, 20 µm.
